# Abnormal behavior is reversible in a chromatin mutant

**DOI:** 10.1101/2023.10.29.564525

**Authors:** Juan D. Rodriguez, Monica N. Reeves, Hsiao-Lin V. Wang, Jaely Z. Chavez, Rhea Rastogi, Sindy R. Chavez, Elicia A Preston, Madhav S. Chadha, Liyang Sun, Emily J. Hill, Victor G. Corces, Karen L. Schmeichel, John I. Murray, David J. Katz

**Author notes:** co-first authors (Juan D. Rodriguez, Monica N. Reeves).

## Abstract

How mutations in histone modifying enzymes lead to neurodevelopmental disorders is unknown. We took advantage of the invariant embryonic lineage and adult nervous system in *C. elegans* to investigate a double mutant between *spr-5/Lsd1/Kdm1a* (H3K4me1/2 demethylase) and *met-2/Setdb1* (H3K9 methyltransferase). We demonstrate that *spr-5; met-2* double mutant worms have a severe chemotaxis defect caused by the ectopic expression of germline genes in somatic tissues. Despite this behavioral defect, we observe few embryonic lineage alterations and an intact adult nervous system. This raises the possibility that the abnormal chemotaxis behavior may be due to ongoing defects in terminally differentiated cells rather than alterations in development. Remarkably, we found that shutting off the ectopic germline expression rescues normal chemotaxis in the same *spr-5; met-2* adult worms that had a chemotaxis defect earlier. This suggests that ongoing inappropriate transcription can block normal behavior in an intact nervous system. Based on these data, it is possible that the intellectual disability and altered behavior observed in human neurodevelopmental syndromes caused by mutations in histone modifying enzymes could be due to ongoing ectopic transcription and may be reversible.

## Introduction

Many human neurodevelopmental disorders are caused by *de novo* mutations in chromatin regulators. For example, Kabuki syndrome is caused by mutations in the H3K4 methyltransferase KMT2D and the H3K27 demethylase KDM6A (1). Kabuki syndrome patients have developmental delay, craniofacial defects and intellectual disability (1). Additionally, four patients have been identified with mutations in the H3K4me1/2 demethylase LSD1/KDM1A (referred to as *spr-5* in *C. elegans*). These LSD1 patients have been referred to as Kabuki-like because their characteristics are nearly identical to Kabuki Syndrome patients (2, 3). However, it remains unclear how defects in the regulation of histone modifications give rise to neurodevelopmental disorders, such as Kabuki Syndrome (4).

H3K4 methylation is acquired co-transcriptionally via the COMPASS complex interacting with RNA polymerase and is associated with actively transcribed genes (5, 6). H3K4 methylation may function as an epigenetic memory, maintaining transcription over time or through mitotic cell divisions (7, 8). As a result, H3K4 methylation that is acquired during the production of gametes may have to be erased to prevent this epigenetic memory from being propagated across generations. In *C. elegans*, SPR-5 is required maternally to erase H3K4me2 and prevent it from being inherited transgenerationally (9). Without SPR-5, worms become increasingly sterile across generations (termed germline mortality) due to the transgenerational accumulation of H3K4me2 and the increasing expression of germline genes. Similarly, when LSD1 is mutated maternally in mice, it results in embryonic lethality at the 2-cell stage (10, 11). This demonstrates that SPR-5/LSD1 has a conserved role in maternal epigenetic reprogramming.

H3K9 methylation is associated with repressed transcription (12). In *C. elegans*, loss of the H3K9 methyltransferase MET-2 (referred to as SETDB1 in mammals) also results in a germline mortality phenotype and *spr-5*; *met-2* double mutants have an exacerbated maternal effect sterility phenotype (13–15). This suggests that SPR-5 and MET-2 cooperate in maternal epigenetic reprogramming. The function of MET-2 in maternal reprogramming is conserved, as maternal loss of SETDB1 in mice also results in early embryonic lethality (16).

During the production of the gametes in *C. elegans*, germline genes acquire the transcription-associated histone modification H3K36me3 via the H3K36 methyltransferase MET-1 (17). In the embryo, this H3K36 methylation is maintained by a transcription-independent H3K36 methyltransferase MES-4 at 176 critical germline genes (17, 18). For the remainder of the manuscript, we will refer to these genes as MES-4 targeted germline genes. At around the 60-cell stage, the germline blastomere P4 divides to give rise to the primordial germ cells, termed Z2 and Z3. Once the embryo hatches, the MES-4 targeted germline genes are expressed as Z2 and Z3 begin to proliferate and loss of MES-4 results in a maternal effect sterility phenotype (17, 18). Thus, it has been suggested that MES-4-dependent H3K36me3 may function as a type of bookmark to help re-specify the germline in the subsequent generation. The transcription factor LSL-1 also plays a critical role in germline function. LSL-1 is first transcribed in P4, continues to be expressed during germline development in all four stages of larval development (L1 to L4), and remains on into the adult. Without, LSL-1 the germline has many defects including defects in meiosis, germline apoptosis and the production of almost no functional gametes (19).

Progeny of *spr-5; met-2* double mutants have a severe developmental delay at the L2 larval stage that is associated with the ectopic expression of MES-4 targeted germline genes in somatic tissues. In addition, progeny of *spr-5; met-2* mutants have ectopic H3K36me3 at MES-4 targeted genes in somatic tissues. When *mes-4* is knocked down by RNA interference in the progeny of *spr-5; met-2* mutants, the ectopic germline expression is eliminated and the L2 larval delay is rescued, suggesting that the severe developmental delay in the progeny of *spr-5; met-2* mutants is caused by the ectopic expression of MES-4 targeted germline genes (20). The ectopic maintenance of H3K36me3 in progeny of *spr-5; met-2* mutants is likely propagated by H3K4 methylation, because the developmental delay of *spr-5; met-2* mutants is also dependent on the H3K4 methyltransferase SET-2 (20). Similar to the progeny of *spr-5; met-2* mutants, loss of a NuRD complex component MEP-1 or DREAM complex component LIN-35 also results in the ectopic expression of germline genes and this ectopic expression is also dependent on MES-4 (21, 22). This suggests that these complexes may be functioning to reinforce SPR-5/MET-2 maternal reprogramming in somatic tissues. In the germline, the function of NuRD is antagonized by the germline transcription factor LSL-1 (19).

In *C. elegans*, the embryonic lineage is completely invariant. This means that every wild-type embryo undergoes the same pattern of cell division and cell migration (23). The invariant cell lineage of *C. elegans* enabled the discovery of programmed cell death (apoptosis) (24). In order to facilitate study of the invariant lineage, the Waterston lab developed automated lineage tracing. The system utilizes a ubiquitously expressed histone-mCherry fusion protein to track cells via their nuclei with 3D time-lapse confocal imaging (25). *C. elegans* also have a simple completely mapped nervous system (26, 27). To enable the study of the nervous system, the Hobert lab developed the NeuroPAL worm in which each neuron expresses a combination of fluorescent molecules that combined with cell position allows the unique identification of all 302 neurons in the adult nervous system of *C. elegans* (28). In the progeny of *spr-5; met-2* double mutants, H3K4me2 is inappropriately inherited from the previous generation and this results in the ectopic expression of MES-4 targeted germline genes (13, 20). Here we take advantage of the invariant embryonic lineage and the simple completely mapped adult nervous system in *C. elegans* to understand how the inappropriate inheritance of chromatin and the ectopic expression of genes in the progeny of *spr-5; met-2* mutants effects individual cells and gives rise to abnormal behavior.

To determine what embryonic cell types the MES-4 targeted germline genes are ectopically expressed in and how this may alter the embryonic lineage, we performed single-cell RNAseq and automated lineage tracing on the progeny of *spr-5; met-2* mutants. We find that MES-4 targeted germline genes begin to be ectopically expressed broadly in many embryonic lineages, except in the germline itself where MES-4 targeted germline genes may not be fully activated. Surprisingly, the ectopic expression of MES-4 targeted germline genes in somatic lineages results in few embryonic lineage defects and does not disrupt the specification of any of the 302 *C. elegans* neurons. This raises the possibility that the ectopic expression of MES-4 germline genes does not primarily alter development, but instead causes an ongoing defect in differentiated cells. While performing these experiments, we noticed that the progeny of *spr-5; met-2* mutants fail to move toward OP50 bacteria. Using chemotaxis assays, we found that the progeny of *spr-5; met-2* mutants have a chemotaxis defect that begins at the L2 stage and is dependent upon MES-4, suggesting that it is dependent on the ectopic expression of MES-4 targeted germline genes. This observation provided the opportunity to test whether the ectopic expression of germline genes may be actively interfering with the function of differentiated cells by determining whether reverting the ectopic expression of these genes restores normal chemotaxis. Remarkably, we find that shutting off the ectopic expression of germline genes after the L2 stage rescues normal chemotaxis behavior. From these data, we conclude that the ectopic expression of germline genes can alter normal behavior in a fully intact nervous system.

## Results

### Germline genes are ectopically expressed in the somatic lineages of *spr-5; met-2* mutant embryos

All of the experiments in this manuscript were performed on F1 progeny of homozygous *spr-5; met-2* double mutants to examine the effect of maternal loss of these enzymes. For simplicity, throughout the manuscript we will refer to these progeny simply as *spr-5; met-2* mutants. Previously we found in *spr-5; met-2* mutants that MES-4 targeted germline genes are ectopically expressed in the soma at the L1 stage (20). To determine how MES-4 targeted germline genes are misexpressed in the embryo at the single-cell level in *spr-5; met-2* mutants, we performed single-cell RNA sequencing on embryos enriched for the 100-200 cell stage. At this stage, early events of gastrulation are completed, zygotic transcription has fully initiated, and most cells have adopted broad tissue identities but are not yet terminally differentiating (29–31). We obtained sequences from 686 *spr-5; met-2* mutant cells and 219 wild-type (N2) cells. To ensure that the transcriptome of our wild-type cells match the published wild-type transcriptome, we integrated our small number of wild-type cells with the published wild-type *C. elegans* single cell atlas (32). To look for potential changes in transcriptomes, we also integrated our *spr-5; met-2* embryonic cells with the published wild-type *C. elegans* single cell atlas (32). Both wild-type and *spr-5; met-2* mutant cells mapped to cell states across the published wild-type dataset (Fig. 1A-C). This indicates that that broad cell classes are not disrupted and cell type specific marker genes are expressed in similar patterns in *spr-5; met-2* mutants. Nevertheless, using a 0.25-fold change cutoff we identified 3,660 genes significantly upregulated and 1,679 genes significantly downregulated in *spr-5; met-2* mutants compared to Wild Type, most of which had relatively low fold changes. The larger number of upregulated genes compared to downregulated genes is consistent with SPR-5 and MET-2 acting as transcriptional repressors.

**Fig. 1.**
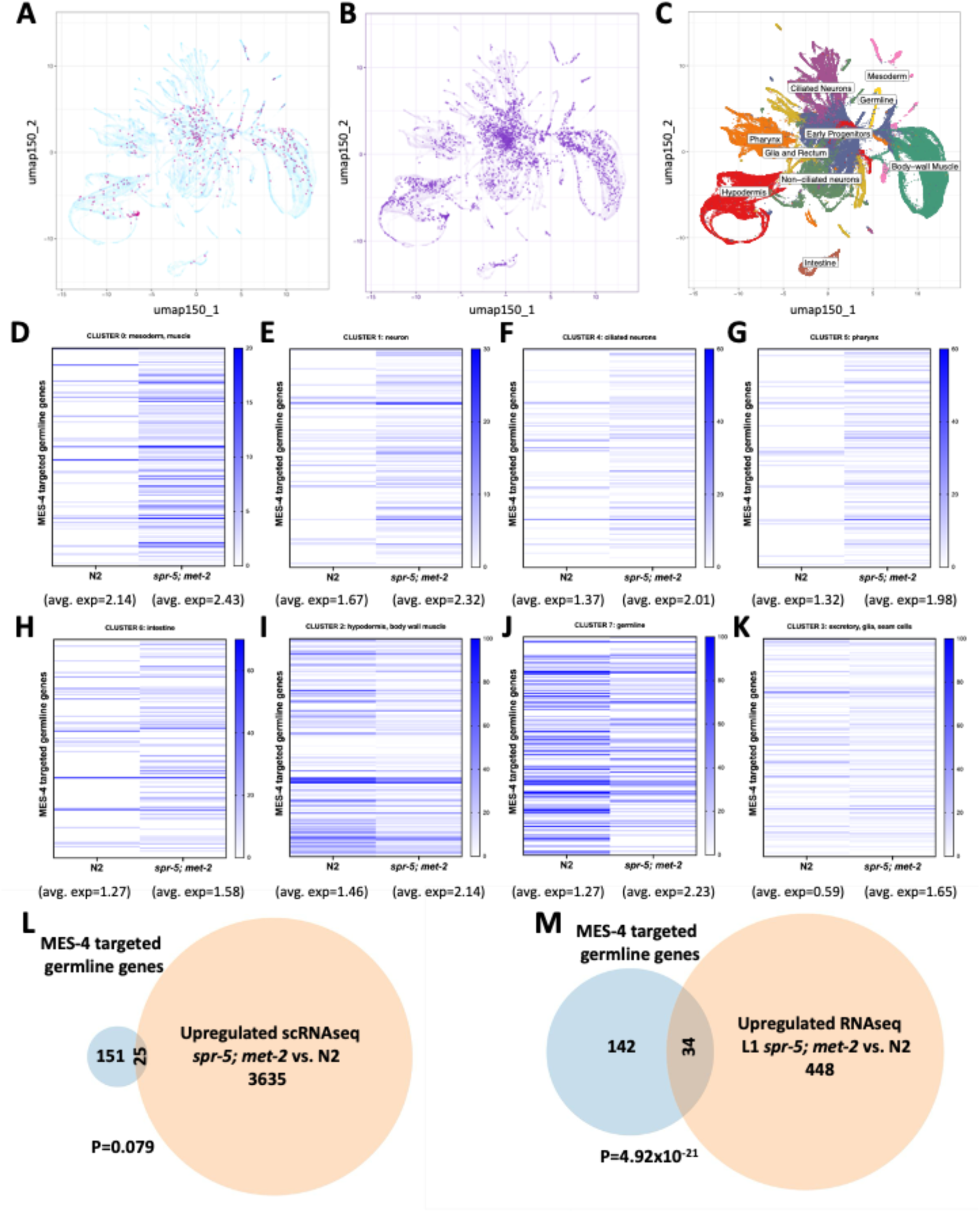
MES-4 targeted germline genes begin to be ectopically expressed during embryogenesis in *spr-5; met-2* mutants. (**A-C**) UMAP projection of all 219 wild-type cells **(A)** and 686 *spr-5; met-2* mutant cells **(B)** from single-cell RNAseq integrated with the published (32) N2 single cell RNAseq data (purple dots show single-cell RNAseq compared to published data in lighter background). The cell type corresponding to each of the published single cell RNAseq clusters is shown in (**C**). (**D-K**) Heat maps corresponding to all 8 clusters obtained from unsupervised hierarchical clustering (from Fig. S1A) showing the percentage of cells that express each of the 197 MES-4 targeted germline genes in Wild Type (N2) compared to *spr-5; met-2* mutants. The average gene expression across all 176 MES-4 targeted germline genes for each cluster is shown below the cluster. (**L,M**) Overlap between MES-4 targeted germline genes and total genes significantly upregulated in *spr-5; met-2* mutants in any of the 8 individual single-cell RNAseq clusters (**L**), or significantly upregulated in *spr-5; met-2* mutants at the L1 larval stage (**M**) from Carpenter et al. 2021 (20). Significance in (**L**) and (**M**) was determined by a hypergeometric test.

To investigate the single cell gene expression data set further, we performed unsupervised hierarchical clustering of our stage-matched wild-type and *spr-5; met-2* mutant cells. Unsupervised clustering identified 8 individual clusters corresponding to major cell classes (germline, muscle, epidermis, etc) and these individual clusters are highly similar between wild-type and *spr-5; met-2* mutants (Fig. S1A,B). This confirms that gene expression is not broadly changed overall in *spr-5; met-2* mutants compared to Wild Type.

To determine whether MES-4 targeted germline genes are ectopically expressed in different lineages, we also examined the gene expression (normalized count matrices) of all MES-4 targeted germline genes in each cluster. The average gene expression across all MES-4 targeted germline genes is higher in *spr-5; met-2* mutants compared to Wild Type (Fig. 1D-K). We also determined the percentage of cells within each cluster that express each of the MES-4 targeted germline genes (Fig. 1D-K). In 5 of the 7 somatic clusters (clusters 0,1,4,5,6) there is a large increase in the percentage of cells within the cluster that express MES-4 targeted germline genes (Fig. 1D-H). Consistent with this, 25 of the 149 examined MES-4 targeted germline genes are significantly ectopically expressed in one of the 7 somatic clusters (Fig. 1L). Together these results suggest that MES-4 targeted germline genes begin to be widely expressed in many somatic lineages in the embryo.

Despite the ectopic expression of MES-4 targeted germline genes in *spr-5; met-2* mutants overall, in the germline cluster (cluster 7) as well as the hypodermis and body wall muscle cluster (cluster 2) the percentage of cells within the cluster that express MES-4 targeted germline genes decreases in *spr-5; met-2* compared to Wild Type (Fig. 1I,J). Consistent with this decrease in the percentage of cells expressing MES-4 targeted genes in the germline, there are only 20 MES-4 targeted germline genes significantly decreased in any cluster and 16 of them are in cells within the germline cluster (cluster 7). However, the decrease in the percentage of cells expressing MES-4 targeted genes in clusters 7 and 2 occurs despite there still being an increase in the average gene expression of the MES-4 targeted germline genes in these clusters (Fig. 1I,J), as there is in all clusters (Fig. 1D-K). Overall, the decrease in the expression of some MES-4 targeted germline genes in the germline cluster (cluster 7) is consistent with the possibility that many MES-4 germline genes fail to activate in the germline of *spr-5; met-2* mutants. It is not clear why MES-4 targeted germline genes are also expressed in fewer cells within cluster 2, but this effect appears to be driven by a higher percentage of these cells expressing MES-4 germline genes in Wild Type. In the excretory, glia and seam somatic cell cluster (cluster 3), the percentage of cells within the cluster that express MES-4 targeted germline genes is unchanged between wild-type and *spr-5; met-2* mutants (Fig. 1K). This raises the possibility that cells of this lineage may be resistant to the ectopic expression of MES-4 targeted germline genes.

To confirm that MES-4 targeted genes are ectopically expressed in *spr-5; met-2* mutants, we performed quantitative RT-PCR on 5 of the MES-4 targeted germline genes that were not significantly misexpressed in our single-cell dataset but were found to be significantly misexpressed at the L1 stage in *spr-5; met-2* mutants in our previous work (20). We found that 4 out of the 5 genes tested were significantly upregulated in 100-200 cell stage embryos with an average fold change of 3.5 (Fig. S1C). Consistent with these results, we previously found by single molecule RNA fluorescence *in situ* hybridization that one of these genes, *htp-1,* is widely ectopically expressed in the somatic cells of the embryo (20). Finally, we also compared the ectopic expression of MES-4 targeted germline genes in our single-cell RNAseq dataset to our previously published *spr-5; met-2* RNAseq performed at the L1 larval stage. In contrast to our single cell dataset, where only 25 MES-4 targeted germline genes are significantly upregulated each only in a single cluster, in our previous L1 RNAseq dataset there were 34 MES-4 targeted genes that were significantly upregulated across the entire L1 larvae (Fig. 1M). This suggests that MES-4 targeted germline genes are increasingly ectopically expressed in larvae compared to embryos. Overall, our single cell RNAseq data suggest that inappropriate chromatin inherited from the previous generation in *spr-5; met-2* mutants is permissive for facilitating ectopic expression beginning widely during embryonic stages, but this ectopic expression increases as development progresses.

### *spr-5; met-2* mutants have a delay in the specification of the germline, but no defects in somatic embryonic development

To determine if cell lineage specification is altered in *spr-5; met-2* mutants, we took advantage of the invariant embryonic lineage in *C. elegans*. By performing automated lineage tracing (33), we tracked the number of cell divisions, timing of cell divisions, cell position and cell migration for all embryonic cells from the 2-cell stage through the 200-cell stage in 10 *spr-5; met-2* mutant embryos, and an additional 8 embryos through the 350-cell stage (Dataset S1,2). Surprisingly, we found no major defects in any somatic lineages deriving from either the AB or P1 blastomeres through the 200-cell stage (∼170 minutes) and only a few significant defects through the 350-cell stage (∼250 minutes) (Fig. 2A-C, Dataset S1,2). The cell division of the pharyngeal intestinal valve cell MSaaapp occurs significantly earlier in *spr-5; met-2* mutants compared to Wild Type (Fig. 2A-C, Fig. S2A,B). In addition, we observe modest delays in the myo-epidermal C lineage and intestinal E lineage in *spr-5; met-2* mutants compared to Wild Type. These defects are less severe than defects seen in mutants for transcription factors involved in fate specification (34–36) and only a few of these cells are significantly delayed (Fig. 2C). Furthermore, the division times of the myo-epidermal C lineage and intestinal E lineage are typically more variable than others even in wild-type animals (36). Despite the slight lineage defects that we observe in *spr-5; met-2* mutants, all embryonic cells are present in the correct place through the 350-cell stage (Fig. 2D,E, Video S1A,B). Overall, these data suggest that somatic development is almost completely normal in *spr-5; met-2* mutant embryos.

**Fig. 2.**
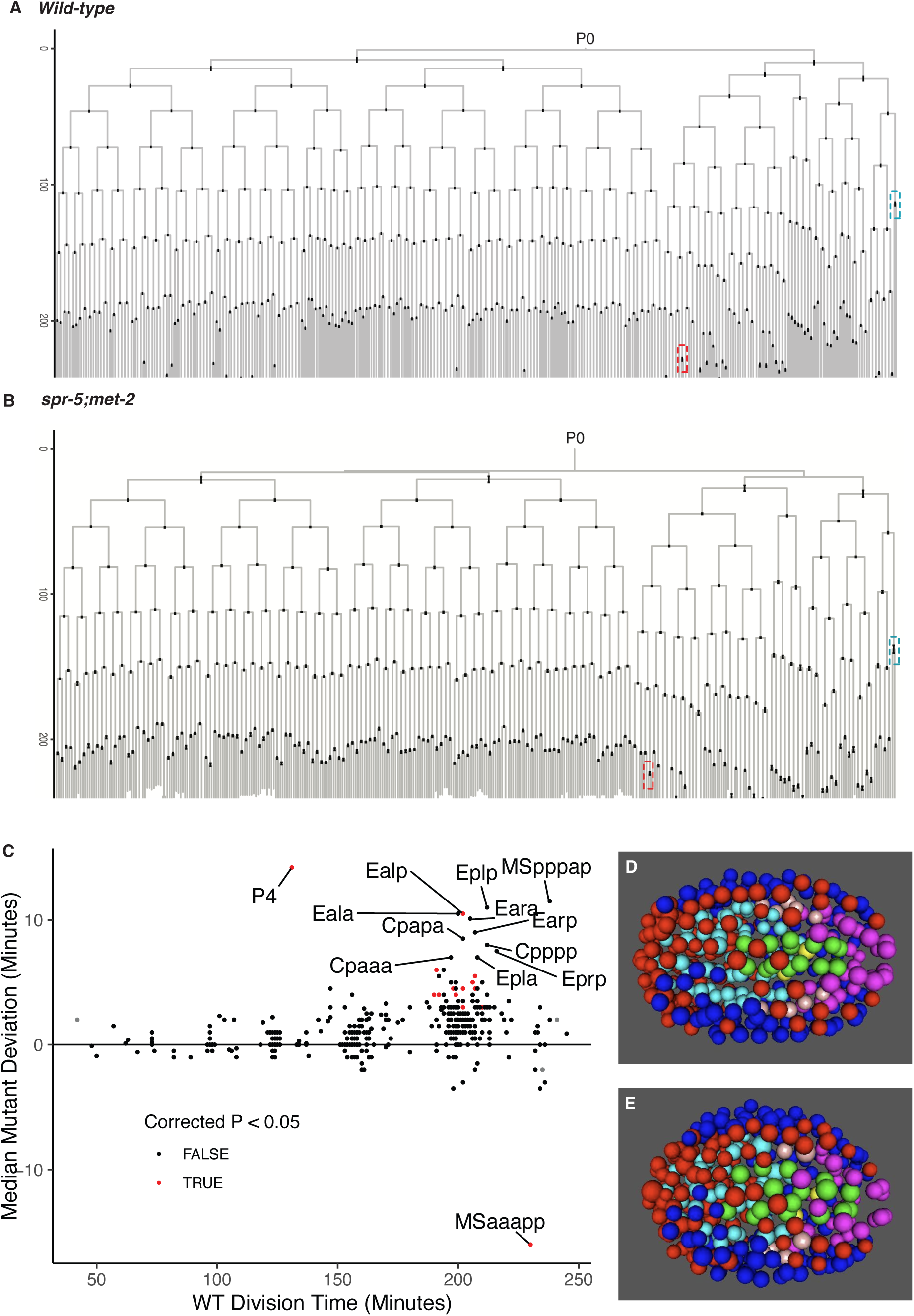
Comparison of the embryonic cell lineage in *spr-5; met-2* versus Wild Type. Wild Type (N2) (**A**) and *spr-5; met-2* (**B**) embryonic lineages through the 350-cell stage. The error bars indicate the SEM from 22 wild-type and 8 *spr-5; met-2* lineages. The Y axis indicates minutes. The red dashed boxes indicate the earlier MSaaapp cell division and the blue dashed boxes indicate the later P_4_ cell division in *spr-5; met-2* compared to Wild Type (See Fig. S2 for detail). (**C**) Plot of the median deviation in minutes of each cell in *spr-5; met-2* (N=18) compared to Wild Type (N=32) from all lineages collected through the 200 and 350 cell stage. Cells that significantly deviate from WT (Bonferroni-corrected P < 0.05; 2 sample t-test, unequal variance) are indicated in red (Dataset S1,2). **(D,E)** Average projection of wild-type **(D)** and *spr-5; met-2* **(E)** embryos at the 350-cell stage (taken from Vid. S1) indicating that all cells are in the correct position. Cells from major lineages are colored as Red:ABa, Blue:ABp, Cyan:MS, Green:E, Purple:C, Pink:D and Yellow:P4.

In contrast, we identified a consistent delay in the germline lineage, with the germline blastomere P4 dividing to give rise to the two primordial germ cells, Z2 and Z3, significantly later in *spr-5; met-2* mutants compared to Wild Type (Fig. 2A-C, Fig S2C,D). The delay in the cell division of the germline blastomere P4 occurs despite there being no delay in the cell D, the somatic sister of P4 (Fig. S2D). The P4 delay correlates with the failure to fully express MES-4 targeted germline genes in the germline cluster (cluster 7)(Fig. 1J), raising the possibility that the delayed division of P4 could be due to the failure to properly activate germline transcription.

### *spr-5; met-2* mutants have a severe defect in chemotaxis towards OP50 bacteria

While maintaining *spr-5; met-2* mutants and performing these experiments, we noticed that *spr-5; met-2* mutants fail to preferentially move towards the OP50 *E. coli* food source. To quantify this behavior defect, we performed a chemotaxis assay and calculated the chemotaxis index (CI). A chemotaxis index of 1.0 indicates that all of the worms are located near the OP50 *E. coli* food source, while a chemotaxis index of 0.0 indicates that the worms are randomly distributed throughout the experimental plate. Compared to wild-type worms which have a chemotaxis index of 0.9, *spr-5* and *met-2* single mutants have a small but significant defect in chemotaxis, with chemotaxis indices of 0.7. In contrast, *spr-5; met-2* mutants have a severe chemotaxis index defect of 0.2, confirming the synergistic interaction between *spr-5* and *met-2* (Fig. 3A, Video S2A). This chemotaxis defect is observed in both L2 larvae and adult animals (Fig. 3). Importantly, in the chemotaxis assay, any animals that fail to move from the origin are not included in the analysis, so the *spr-5; met-2* chemotaxis defect is not due to a lack of motility. Taken together, these results demonstrate that *spr-5; met-2* chemotaxis mutants fail to preferentially go towards the OP50 *E. coli* food source, despite embryonic development being almost completely normal in these mutants.

**Fig. 3.**
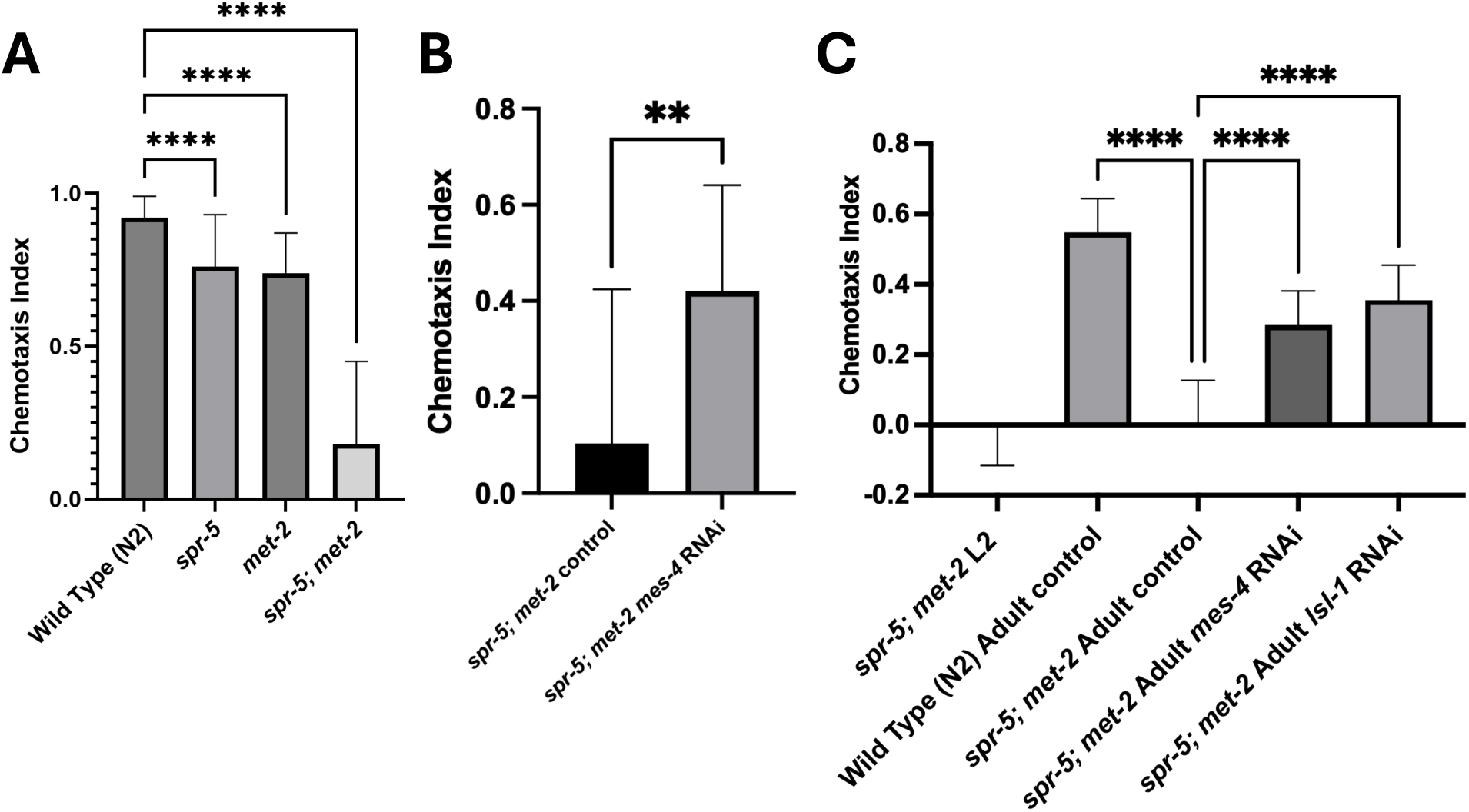
Blocking the ectopic transcription of MES-4 targeted germline genes in L2 *spr-5; met-2* mutants that already have a chemotaxis defect restores normal chemotaxis. (**A**) The chemotaxis index of Wild Type (N2) (N=1008) (Video S2A), *spr-5* (N=1661), *met-2* (N=1117) and *spr-5; met-2* (N=263) adults. (**B**) The chemotaxis index of *spr-5; met-2* mutant adults is rescued when *spr-5; met-2* mutants are developed in the presence of *mes-4* RNAi (N= 948) compared to control (L4440) RNAi (N=690). (**C**) The chemotaxis index of L2 *spr-5; met-2* double mutants (N=2950) (Video S2B) is significantly rescued by shifting on to *mes-4* (N=763) (Video S2C) or *lsl-1* (N=776) RNAi (Video S2E) at the L2 stage and then assaying chemotaxis in the resulting adults. These resulting adults, which are the same worms that previously had a chemotaxis defect at the L2 stage, are rescued to levels that are similar to Wild Type (N2) (N=1003) under these conditions. In contrast, control (L4440) RNAi does not rescue (N=603) (Video S2D). Worms were assayed as adults ∼5 days after initiating RNAi at the L2 stage. The chemotaxis in **B** and **C** was performed with OP50 bacteria after carrying out RNAi using HT115 bacteria. The error bars represent the S.E.M.. Significance was calculated in (**A,C**) by one-way ANOVA and (**B**) by unpaired t-test. ****≤0.0001, **≤0.01.

### The *spr-5; met-2* chemotaxis defect is dependent upon the ectopic expression of MES-4 targeted germline genes

Previously we demonstrated that a severe larval (L1/L2 stage) developmental delay also observed in *spr-5; met-2* mutants can be rescued by knockdown of the H3K36 methyltransferase MES-4 (20). This suggested that the developmental delay is dependent upon the ectopic expression of MES-4 targeted germline genes. To determine whether the chemotaxis defect of *spr-5; met-2* mutants is also dependent upon the ectopic expression of MES-4 germline genes, we measured chemotaxis in *spr-5; met-2* mutants which developed in the presence of *mes-4* RNA interference (RNAi). RNAi of *mes-4* in *spr-5; met-2* mutants significantly rescued the chemotaxis defect, suggesting that the chemotaxis defect is also dependent upon the ectopic transcription of MES-4 targeted germline genes (Fig. 3B).

### Zygotic knockdown of the ectopic transcription of MES-4 targeted germline genes in L2 *spr-5; met-2* mutants restores normal chemotaxis in these same animals as adults

The lack of embryonic developmental defects in *spr-5; met-2* mutants raised the possibility that the severe chemotaxis defect in these mutants could be due to the ongoing ectopic expression of MES-4 targeted germline genes. If this is the case, it might be possible to shut off the ectopic expression of the MES-4 targeted germline genes and restore normal chemotaxis. To test this possibility, we took L2 *spr-5; met-2* larvae which already have a severe chemotaxis defect and attempted to shut off the ectopic expression of MES-4 targeted germline genes by shifting to *mes-4* RNAi at the L2 stage, and then assaying chemotaxis in the resulting adults. Importantly, because *spr-5; met-2* double mutants have a severe developmental delay which is not rescued when *mes-4* RNAi is initiated at the L2 stage, it takes ∼5 days for them to develop from L2 larvae to adults. This gives the RNAi plenty of time to knockdown the ectopic transcription of the MES-4 targeted germline genes in somatic tissues. It is also possible that RNAi works better in the soma of *spr-5; met-2* double mutants because of the ectopic expression of germline RNAi machinery (20). In these experiments (and the previous MES-4 rescue experiments in Fig. 3B) the RNAi was performed with HT115 bacteria rather than OP50, prior to assaying chemotaxis towards the OP50 *E. coli* food source. This results in an overall reduction in the chemotaxis index (Fig. 3C). For example, under these conditions the chemotaxis index of wild-type animals is 0.55 (or 0.4 in the previous MES-4 rescue experiments in Fig. 3B) rather than 0.9, and the chemotaxis index of *spr-5; met-2* double mutants is 0.0 (or 0.1 in the previous MES-4 rescue experiments in Fig. 3B) rather than 0.2.

Remarkably, we find that shutting off the ectopic germline expression of the MES-4 targeted germline genes after the L2 stage results in a significant rescue of the chemotaxis index from 0.0 to 0.28 in these same animals at the adult stage (Fig. 3C, Video S2B-D). MES-4 is required to propagate the ectopic transcription of MES-4 targeted germline genes to somatic tissues but may not be completely required to continually maintain the ectopic transcription of MES-4 targeted germline genes in somatic cells. Therefore, we also tried to shut off the ectopic expression of germline genes by performing RNAi against *lsl-1.* LSL-1 is a germline transcription factor that is actively required for the transcription of many germline expressed genes, including some MES-4 targeted germline genes (19). We found that knockdown of LSL-1 after the L2 stage in *spr-5; met-2* mutants results in a stronger rescue of the chemotaxis index in *spr-5; met-2* mutants from 0.0 to 0.36 (compared to 0.28 with *mes-4* RNAi), such that the chemotaxis index is similar to Wild Type (0.55) under these experimental conditions (Fig. 3C, Video S2E). As with the MES-4 RNAi experiments, these chemotaxis experiments were performed in the same adult animals that were previously defective at the L2 stage. Our finding that the chemotaxis behavior is reversible by shutting off the ectopic expression of germline genes is even more remarkable considering that the nervous system is already formed in L2 larvae when we begin the RNAi. Taken together these results suggest that the ongoing ectopic transcription of MES-4 targeted germline genes can block normal chemotaxis behavior in a fully intact nervous system.

### The adult nervous system is completely present in *spr-5; met-2* double mutants

Our finding that knockdown of *mes-4* or *lsl-1* by RNAi rescues normal chemotaxis in *spr-5; met-2* double mutants suggests that the nervous system is fully intact in these animals. However, to verify that all of the neurons are present in *spr-5; met-2* double mutants, we took advantage of the invariant adult nervous system in *C. elegans*, which can be imaged using the NeuroPAL system (28). To verify that the NeuroPAL system is working, we first counted all 302 unique neurons in the wild-type NeuroPAL strain (Fig. 4A,B,E). Identical to the wild-type animals, we also counted 302 unique neurons in *spr-5; met-2* mutants (Fig. 4C,D,E). This includes the key neurons that are known to specifically function in chemotaxis towards the OP50 *E. coli* food source: the left/right pairs of sensory neurons AWA and AWC, the left/right pairs of interneurons AIA, AIB and AIY, as well as the left/right pair of motor neurons RIM (Video S3A,B) (37–43). These data confirm that all neurons are present in adult *spr-5; met-2* double mutants. Thus, in *spr-5; met-2* mutants the ongoing ectopic transcription of MES-4 targeted germline genes can block normal chemotaxis behavior in a fully intact nervous system, in a way that is reversible by shutting off the ectopic transcription.

**Fig. 4.**
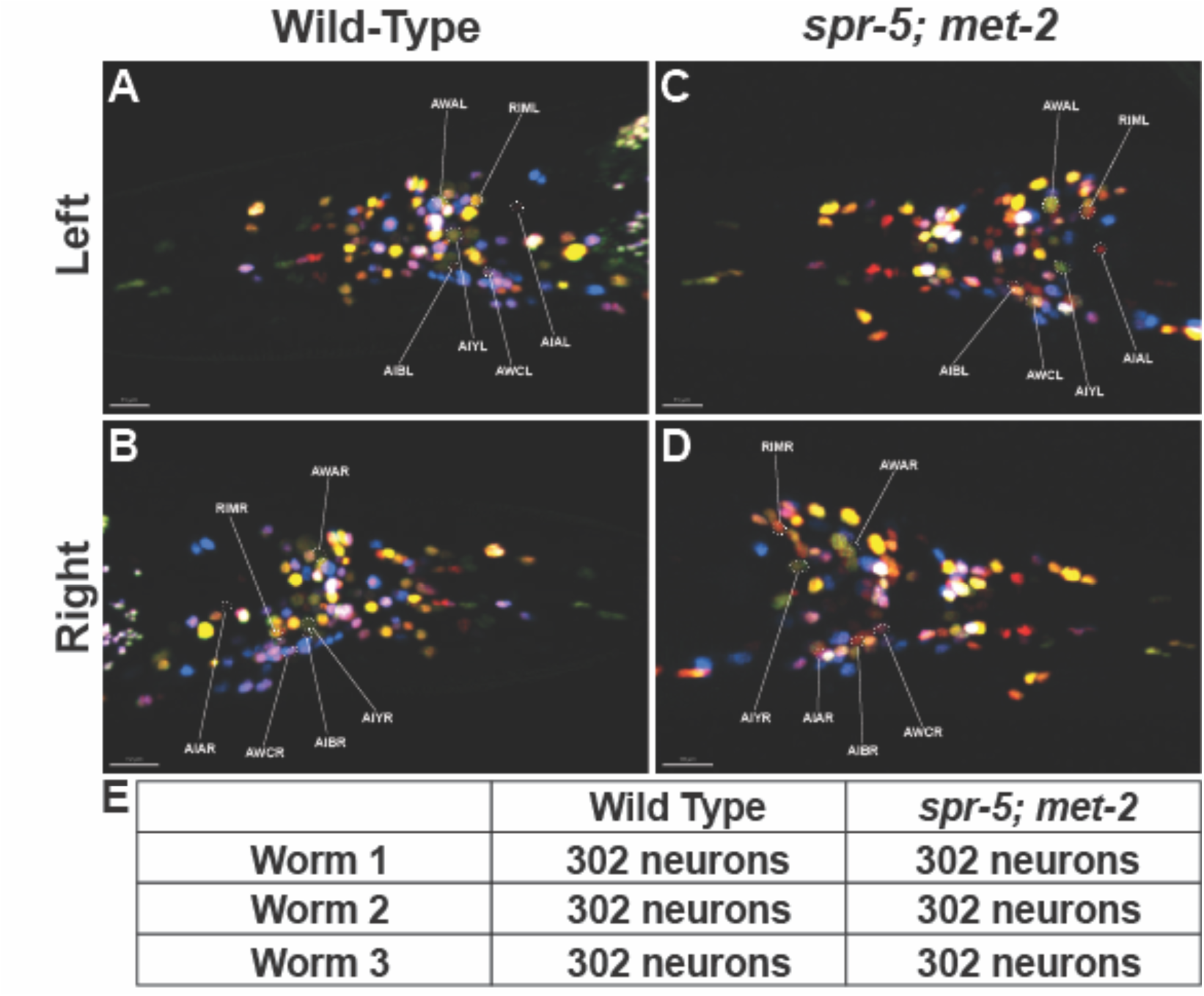
The nervous system is completely intact in *spr-5; met-2* mutants with impaired chemotaxis. Example left side (**A**,**C**) and right side (**B**,**D**) projections of wild-type (**A**,**B**) and *spr-5; met-2* mutant (**C**,**D**) adult heads. The left/right pairs of sensory neurons (AWA and AWC) and interneurons (AIA, AIB and AIY), as well as the left/right pair of motor neurons RIM that are known to specifically function in chemotaxis towards the OP50 *E. coli* food source are labelled (3D rotation in Vid. S3A,B). (**E**) Table listing the number of uniquely identified neurons in wild-type and *spr-5; met-2* mutant worms (N=4 biological replicates).

## DISCUSSION

*spr-5; met-2* double mutants inappropriately inherit chromatin from the previous generation and ectopically express germline genes in somatic tissues. This provided the unique opportunity to determine how the ectopic expression of genes resulting from inappropriately inherited histone methylation influences behavior. Previously, we found that *spr-5; met-2* mutants have a severe L2 developmental delay that is associated with the ectopic expression of MES-4 targeted germline genes. Knock down of *mes-4* by RNA interference eliminates the ectopic expression of MES-4 targeted germline genes and rescues the L2 developmental delay, suggesting that the developmental delay is dependent upon the ectopic expression of MES-4 targeted germline genes (20). Here we found that *spr-5; met-2* mutants also have a severe defect in chemotaxis towards OP50 bacteria at both the L2 larval stage and in adults. There is also a significant chemotaxis defect in both *spr-5* and *met-2* single mutants, but the chemotaxis defect is much more severe in *spr-5; met-2* double mutants, confirming the synergistic interaction between *spr-5* and *met-2*. In addition, we find that the severe chemotaxis defect in *spr-5; met-2* mutants is dependent upon MES-4, suggesting that this phenotype is also dependent upon the ectopic expression of MES-4 targeted germline genes.

Single-cell RNAseq analysis of *spr-5; met-2* mutants at the 100-200 cell stage demonstrated the MES-4 targeted germline genes begin to be ectopically expressed in the embryo. However, the ectopic expression was less severe than what we previously observed at the L1 stage (20). One potential explanation for the increased ectopic expression at the L1 stage is that the permissive chromatin must be acted upon by transcription factors that are not expressed until the individual lineages are further differentiated.

The ectopic expression of MES-4 targeted germline genes is observed equally in most somatic clusters. This is consistent with a model in which the ectopic chromatin is inherited fairly uniformly across embryonic lineages. However, we do not observe the ectopic expression of the MES-4 targeted germline genes in the excretory, glia and seam somatic cell cluster (3), so some lineages may not inherit the ectopic chromatin or may be resistant to the ectopic expression of these germline genes. Despite the ectopic expression of MES-4 germline genes, unsupervised clustering analysis of our *spr-5; met-2* single-cell RNAseq data demonstrated that these mutants are very similar to Wild Type. This suggests that the ectopic transcription does not broadly interfere with cell specification in the embryo. Consistent with this possibility, our automated lineage tracing of *spr-5; met-2* mutants demonstrated that there are very few defects in the embryonic lineage. Using Neuropal imaging, we also observed that *spr-5; met-2* mutants have all 302 adult neurons, including all the neurons that are known to be required for normal chemotaxis towards the OP50 *E. coli* food source. The lack of embryonic lineage and adult nervous system defects is surprising because they occur despite there being a severe developmental delay and impaired chemotaxis beginning at the L2 stage.

The lack of defects caused by the inappropriate somatic expression of germline genes is consistent with two possible models. One possibility is that embryonic development is resistant to the MES-4 targeted germline genes that are significantly ectopically expressed in the embryo. In this scenario, development may overcome the ectopic expression. Alternatively, it is possible that the critical MES-4 targeted germline genes that cause the L2 developmental delay and chemotaxis defect are not yet sufficiently misexpressed in the wrong cell types or expressed at a sufficient level to cause these defects in the embryo. Regardless, the lack of somatic embryonic lineage defects and the intact nervous system raised the possibility that the developmental delay and chemotaxis defects in *spr-5; met-2* mutants are not due to a failure to properly specify a specific lineage during embryogenesis.

Although we do not observe any somatic defects in the embryonic lineage of *spr-5; met-2* mutants, we detect a significant delay in duration of the germline blastomere P4 before dividing to give rise to the primordial germ cells Z2 and Z3 in *spr-5; met-2* mutants compared to Wild Type. This delay occurs despite there being no corresponding delay in the P4 sister cell D, arguing that the P4 delay is highly specific to the germline lineage. While it remains unclear why there is a delay in the P4 cell division, our single-cell data demonstrate that MES-4 targeted genes fail to be fully expressed in the germline cluster (7). Therefore, it is possible that the failure to activate the transcription of the MES-4 germline genes causes a delay in the P4 cell division. The failure to fully activate germline transcription in Z2 and Z3 may also account for why *spr-5; met-2* mutants are completely sterile (13, 14). Previously, it has been shown that the target of SPR-5, H3K4me2, is specifically lost during the P4 cell division to generate Z2 and Z3. Thus, it is possible that an increase in H3K4me2 in *spr-5; met-2* mutants results in a failure to erase H3K4me2 in Z2 and Z3. This in turn could lead to the delay in the P4 cell division, but this possibility remains to be examined.

The lack of somatic embryonic lineage defects in *spr-5; met-2* mutants raised the possibility that the chemotaxis defect is due to the ongoing ectopic expression of germline genes. If this were the case, we would expect that eliminating the ectopic expression in worms that already have a chemotaxis defect would eliminate the chemotaxis defect. To test this possibility, we performed *mes-4* RNAi beginning at the L2 stage in *spr-5; met-2* mutants to block the ectopic expression of MES-4 targeted germline genes, when these mutants already have a chemotaxis defect. Importantly, at the L2 larval stage *C. elegans* already have a completely intact nervous system. Remarkably, we find that knock down of *mes-4* beginning at the L2 stage significantly rescues normal chemotaxis behavior in the same animals that were previously defective at the L2 stage. MES-4 is thought to function in propagating the inappropriate expression of MES-4 targeted germline genes in somatic tissues and may not be completely required to maintain the ectopic expression of these genes. Therefore, we also tried to block the ectopic expression of germline genes by performing RNA interference against the germline transcription factor LSL-1. LSL-1 is required for the transcription of many germline genes, including many MES-4 targeted germline genes. Knock down of *lsl-1* beginning at the L2 stage in *spr-5; met-2* mutants resulted in an even stronger rescue of normal chemotaxis behavior than knock down of *mes-4.* Taken together these results suggest that the ectopic expression of MES-4 targeted germline genes actively interferes with normal chemotaxis behavior in a completely intact nervous system.

It is unclear how the ectopic expression of germline genes actively interferes with normal chemotaxis behavior. One possibility is that some normal component of neuronal function is blocked at the transcription level. For example, the expression of synaptic proteins or a chemoreceptor could be actively inhibited in *spr-5; met-2* mutants. An alternative possibility that is not mutually exclusive is that some aspect of germline function actively interferes with neuronal function. For example, activation of meiosis genes could cause inappropriate chromosomal condensation in neurons. Regardless, our finding that ectopic transcription actively interferes with an intact nervous system has implications for neurodevelopmental diseases, such as Kabuki Syndrome and LSD1 patients. Based on our results, it is possible that the intellectual disability or altered behavior in these patients could be due to an ongoing defect in a properly formed nervous system. Even though these patients have *de novo* mutations that were either inherited from one of their parents or mutated in the very early embryo, it is possible that the inappropriately inherited chromatin results in a defect that only manifests itself later in development because the transcription factors that are required to interact with the permissive chromatin are only activated in certain differentiated cell types, such as neurons. Regardless of the etiology, if the nervous system defects in these patients are due to the ongoing ectopic expression of genes, it may be possible to rescue these defects by turning off the ectopic transcription. Consistent with this possibility, it has been recently shown that behavioral defects in two mouse models of epigenetic syndromes can be rescued at the adult stage. Loss of the DNA methylation binding protein MeCP2 that causes Rhett Syndrome can be rescued by re-expressing MeCP2 in adult mice (44–46). In addition, heterozygous loss of the H3K4 methyltransferase *Kmt2d* that causes Kabuki Syndrome can be partially rescued in adult mice with a histone deacetylase inhibitor (47). Our data provide a possible explanation for these results.

## Materials and Methods

### Strains

All *C. elegans* strains were cultured at 20°C on 60 mm nematode growth media (NGM) agar plates with OP50 bacteria grown in Luria Broth (LB). Strains used were: N2: Wild Type (Bristol isolate); the *C. elegans spr-5 (by101)(I)* strain was provided by R. Baumeister (Albert Ludwig University of Freiburg, Germany); the MT13293: *met-2 (n4256)(III) strain* was provided by R. Horvitz (Massachusetts Institute of Technology, MA, USA); JIM113: ujIs113 [Ppie-1::H2B::mCherry, unc-119(+); Pnhr-2::HIS-24::mCherry, unc-119(+) II. *spr-5(b101)/tmC27[unc-75(tmls1239)]* I*; met-2(n4256)]/qC1 qIs26 [lag-2::GFP + pRF4 rol-6(su1006)]* III strain was created for maintenance as a double heterozygote. The automated cell lineage strain is *spr-5(b101)/tmC27[unc-75(tmls1239)]* I; JIM113: ujIs113 [Ppie-1::H2B::mCherry, unc-119(+); Pnhr-2::HIS-24::mCherry, unc-119(+) II; *met-2(n4256)]/qC1 qIs26 [lag-2::GFP + pRF4 rol-6(su1006)]* III. The NeuroPAL strain otIs669 V (Wild Type) was obtained from *Caenorhabditis* Genetics Center (CGC) and crossed into *spr-5; met-2* mutants.

### Single-worm genotyping

Single animals were picked into 10-12 μl of lysis buffer (50 mM KCl, 10 mM Tris-HCl (pH 8.3), 2.5 mM MgCl_2_, 0.45% NP-40, 0.45% Tween-20, 0.01% gelatin) containing 1 µg/µl final concentration of proteinase K and incubated at −80°C overnight followed by 60°C for 1 hour and then 95°C for 15 minutes. PCR reactions were performed with AmpliTaq Gold (Invitrogen) according to the manufacturer’s protocol and reactions were resolved on agarose gels. The following genotyping primers were used: spr-5 (*by101*)I: fwd: AAACACGTGGCTCCATGAAT, rev(wt):GAGGTTTTGAGGGGTTCCAT, rev(mut):CTTGAAACAGACTTGAACATCAAAGATCGG; met-2 (n4256): fwd(wt):GTCACATCACCTGCATCAGC, rev(wt):ATTTCATTACGGC TGCCAAC, fwd(mut):ATTCGAAAAATGGACCGTTG, rev(mut):TCTATTCCCAGGAGCCAATG;

### Quantitative PCR

Total RNA was isolated using TRIzol reagent (Invitrogen) from synchronized embryos at room temperature. cDNA synthesis and qPCR were carried out as previously described (13). mRNA was quantified by real-time PCR, using iO SYBR Green Supermix (BioRad). The following primers were used: *htp-1* (ATTCGGAGGACAGTGACACAA and GTGCTTTCTCGAGAGACTCAGTTATATC) *cpb-1* (GTGCTGATTGATTGGCCTCG and CCGTTACAGCGCGTGAACCG); *rmh-1* (TGTAGTCATTATGCCAAGTATCTGC and ATCTGTTACTCGTATCTGTAGTAGCC); *ftr-1* (TCCGCTCACTTCGAATACGG and TACCATCGCGATTGTGAGC); *fbxa-101* (TATCGAAGACAAGCTCGCCG and TGCGAACGGAAATCCAATCG); *ama*-1 (TACCTACACTCCAAGTCCATCG and CGATGTTGGAGAGTACTGAG). Each mRNA expression was normalized to *ama-1* control. Fold enrichment was calculated as mutant/Wild Type (N2).

### RNA interference

*Escherichia coli* HT115 transformed with a vector expressing dsRNA of *mes-4*, *lsl-1*, L4440 or *ama-1* obtained from the Ahringer library (Source BioScience). RNAi bacteria and empty vector control were grown at 37°C and seeded on RNAi plates (standard NGM plates containing ampicillin at 100 mg/ml, tetracycline at 5 mg/ml and isopropylthiogalactoside (IPTG) at 0.4 mM. RNAi plates were left at room temperature to induce for at least 48 hr.

### Chemotaxis assay

The experimental 60mm NGM plate was divided into four equal quadrants with 5µL of control LB/vehicle in two quadrants (C1 and C2) and 5µL of 200X concentrated fresh overnight cultures of *Escherichia coli* OP50 in the other two quadrants (E1 and E2). The plate was placed into the sterile hood for drying purposes. Worms were collected and rinsed 3 times with M9 buffer (22 mM KH_2_PO_4_, 42 mM Na_2_HPO_4_, 86 mM NaCl, and 1 mM MgSO_4_). A total of ∼50 worms per plate were placed into the center of the experimental plate. The plates were then incubated at 20°C for 1 hr. Worms that moved to any of the quadrants after the incubation time were recorded. The Chemotaxis Index was calculated as follows; (E1 + E2) – (C1 + C2)/ Total worms moved.

### Lineage tracing

Cell lineage tracing was carried out as previously described using a Zeiss LSM 510 confocal microscope with some minor changes (33). While using the 20µm beads, we noticed that the embryo was moving out of focus. To rectify this, an NGM plate was used as a platform to hold the embryo. This alternative approach is described in detail (48). Additional embryos were imaged using a Leica Stellaris confocal microscope. Cell lineages were traced using StarryNite, curated using AceTree, and Cell positions and division times compared to a wild-type compendium as in (25, 33, 35, 36, 49–51).

### Single-cell RNA sequencing and data analysis

Single-cell isolation was performed according to Packer et al. (32) with minor modifications. For the 100-200 cell stage, only one enzyme (Chitinase) was used for embryo eggshell disruption and single cells dissociation (48). Single-cell RNA sequencing was performed using the 10X Genomics single-cell capturing system. Single cells were isolated from embryos produced by approximately 1,000 hand-selected *spr-5; met-2* mutant mother worms per strain. 1,000 to 3,000 cells were loaded on the 10X Genomics Chromium Controller. Single-cell cDNA libraries were prepared using the Chromium Next GEM Single Cell 3ʹ LT Reagent Kits v3.1 (Dual Index). Libraries were sequenced by FSU College of Medicine Translational Science Laboratory (Florida State University, FL, USA). After the 10X QC, N2 had 219 estimated cells, 62,491 mean reads per cell and 1,067 median genes per cell. *spr-5; met-2* had 686 cells, 20,763 mean reads per cell and 526 median genes per cell. The number of cells was limited by the need to hand-pick *spr-5; met-2* double mutant mothers prior to embryo collection. Sequencing: 28 million paired-end of ∼150 bp paired-end reads (Illumina NextSeq 600). The Cell Ranger Software Suite 3.0.2 (10X Genomics) was used for the alignment of the single-cell RNA-seq output reads and generation of feature, barcode and matrices. Seurat analysis was used for unsupervised hierarchical cluster formation.

### NeuroPAL

10-15 young adult stage worms were paralyzed with 10mM levamisole and mounted on a 1.5% agarose gel pad. The coverslip was sealed with nail polish to reduce evaporation and warping of the gel pad during lengthy imaging. Worms were imaged on a Nikon A1R HD25 inverted confocal microscope with optical configurations and imaging channels from the NeuroPAL manual “Configuring Your Microscope for NeuroPAL v3,” obtained from Hobert Lab website https://www.hobertlab.org/neuropal/ (28). Images were taken at 60X with 4 channels: CyOFP, pseudocolored green, mNeptune2.5, pseudocolored red, mTagBFP2, pseudocolored blue, and TagRFP-T panneuronal marker, pseudocolored white, using a Z-stack with 30-50 steps of 0.8μm. A minimum of 3 images were taken of each worm: head, midbody, and tail. The resulting images were adjusted for brightness/contrast and channels were merged into a composite using Fiji/ImageJ (52). 3D images were obtained using Imaris software version 9.8.0 as described in (28, 53). Imaris 3D images were used to count the number of neurons in each worm and annotate the chemotactic circuit neurons. Neurons were identified using the “NeuroPAL Reference Manual v1” and “Using NeuroPAL for ID v1” manuals obtained from Hobert Lab website.

## Data, Materials and Software Availability

The single cell RNAseq data has been deposited in GEO under accession number GSE272897.

## Acknowledgements

First and foremost, we thank S. Brenner for the foresight to develop a system (*C.* elegans) that is perfect for addressing this question. We are grateful to members of the Katz lab, T. Caspary and C. Bean for their helpful discussion; C. Huynh, and A. Santella for advice on lineage tracing and single cell RNAseq; the Emory University Emory Integrated Cellular Imaging Core Facility (RRID:SCR_023534) for help and support with imaging. We thank R. Horvitz and the *Caenorhabditis* Genetics Center (funded by NIH P40 OD010440) for strains and WormBase (WS292). Juan D. Rodriguez was supported by NIH National Institute of Child Health and Human Development 3F31HD100145-03S1 grant and the training grant (T32GM008490-21). Monica Reeves was supported by training grants (T32GM008490-28) and (T32GM149422-01). This work was funded by a grant to D.J.K. (NSF IOS1354998) and to J.I.M (NIH R35GM127093 and R35GM153497).

## References

1. M. P. Adam et al., Kabuki syndrome: international consensus diagnostic criteria. J Med Genet 56, 89–95 (2019).

2. J. X. Chong et al., Gene discovery for Mendelian conditions via social networking: de novo variants in KDM1A cause developmental delay and distinctive facial features. Genet Med 18, 788–795 (2016).

3. M. Melo, J. Oliveira, D. Antunes, eP189: Insights into the phenotype of KDM1A-related neurodevelopmental disorder: A new chromatinopathy. Genetics in Medicine 24 (2022).

4. M. Gabriele, A. Lopez Tobon, G. D’Agostino, G. Testa, The chromatin basis of neurodevelopmental disorders: Rethinking dysfunction along the molecular and temporal axes. Prog Neuropsychopharmacol Biol Psychiatry 84, 306–327 (2018).

5. N. J. Krogan et al., Methylation of histone H3 by Set2 in Saccharomyces cerevisiae is linked to transcriptional elongation by RNA polymerase II. Mol Cell Biol 23, 4207–4218 (2003).

6. B. K. Cenik, A. Shilatifard, COMPASS and SWI/SNF complexes in development and disease. Nat Rev Genet 22, 38–58 (2021).

7. B. Li, M. Carey, J. L. Workman, The role of chromatin during transcription. Cell 128, 707–719 (2007).

8. T. W. Lee, D. J. Katz, Hansel, Gretel, and the Consequences of Failing to Remove Histone Methylation Breadcrumbs. Trends Genet 36, 160–176 (2020).

9. D. J. Katz, T. M. Edwards, V. Reinke, W. G. Kelly, A C. elegans LSD1 demethylase contributes to germline immortality by reprogramming epigenetic memory. Cell 137, 308–320 (2009).

10. J. A. Wasson et al., Maternally provided LSD1/KDM1A enables the maternal-to-zygotic transition and prevents defects that manifest postnatally. Elife 5 (2016).

11. K. Ancelin et al., Maternal LSD1/KDM1A is an essential regulator of chromatin and transcription landscapes during zygotic genome activation. Elife 5 (2016).

12. J. Padeken, S. P. Methot, S. M. Gasser, Establishment of H3K9-methylated heterochromatin and its functions in tissue differentiation and maintenance. Nat Rev Mol Cell Biol 23, 623–640 (2022).

13. S. C. Kerr, C. C. Ruppersburg, J. W. Francis, D. J. Katz, SPR-5 and MET-2 function cooperatively to reestablish an epigenetic ground state during passage through the germ line. Proc Natl Acad Sci U S A 111, 9509–9514 (2014).

14. E. L. Greer et al., A Histone Methylation Network Regulates Transgenerational Epigenetic Memory in C. elegans. Cell Rep S2211-1247(14)00158-2 [pii]10.1016/j.celrep.2014.02.044 (2014).

15. E. C. Andersen, H. R. Horvitz, Two C. elegans histone methyltransferases repress lin-3 EGF transcription to inhibit vulval development. Development 134, 2991–2999 (2007).

16. J. Kim et al., Maternal Setdb1 Is Required for Meiotic Progression and Preimplantation Development in Mouse. PLoS Genet 12, e1005970 (2016).

17. A. Rechtsteiner et al., The histone H3K36 methyltransferase MES-4 acts epigenetically to transmit the memory of germline gene expression to progeny. PLoS Genet 6 (2010).

18. H. Furuhashi et al., Trans-generational epigenetic regulation of C. elegans primordial germ cells. Epigenetics Chromatin 3, 15 (2010).

19. D. Rodriguez-Crespo, M. Nanchen, S. Rajopadhye, C. Wicky, The zinc-finger transcription factor LSL-1 is a major regulator of the germline transcriptional program in Caenorhabditis elegans. Genetics 221 (2022).

20. B. S. Carpenter et al., Caenorhabditis elegans establishes germline versus soma by balancing inherited histone methylation. Development 148 (2021).

21. Y. Unhavaithaya et al., MEP-1 and a homolog of the NURD complex component Mi-2 act together to maintain germline-soma distinctions in C. elegans. Cell 111, 991–1002 (2002).

22. L. N. Petrella et al., synMuv B proteins antagonize germline fate in the intestine and ensure C. elegans survival. Development 138, 1069–1079 (2011).

23. J. E. Sulston, E. Schierenberg, J. G. White, J. N. Thomson, The embryonic cell lineage of the nematode Caenorhabditis elegans. Dev Biol 100, 64–119 (1983).

24. J. E. Sulston, H. R. Horvitz, Post-embryonic cell lineages of the nematode, Caenorhabditis elegans. Dev Biol 56, 110–156 (1977).

25. Z. Bao et al., Automated cell lineage tracing in Caenorhabditis elegans. Proc Natl Acad Sci U S A 103, 2707–2712 (2006).

26. J. G. White, E. Southgate, J. N. Thomson, S. Brenner, The structure of the nervous system of the nematode Caenorhabditis elegans. Philos Trans R Soc Lond B Biol Sci 314, 1–340 (1986).

27. S. J. Cook et al., Whole-animal connectomes of both Caenorhabditis elegans sexes. Nature 571, 63–71 (2019).

28. E. Yemini et al., NeuroPAL: A Multicolor Atlas for Whole-Brain Neuronal Identification in C. elegans. Cell 184, 272–288 e211 (2021).

29. S. M. Robertson, P. Shetty, R. Lin, Identification of lineage-specific zygotic transcripts in early Caenorhabditis elegans embryos. Dev Biol 276, 493–507 (2004).

30. L. R. Baugh, A. A. Hill, D. K. Slonim, E. L. Brown, C. P. Hunter, Composition and dynamics of the Caenorhabditis elegans early embryonic transcriptome. Development 130, 889–900 (2003).

31. E. A. Bucher, G. Seydoux, Gastrulation in the nematode Caenorhabditis elegans. Seminars in Developmental Biology 5, 121–130 (1994).

32. J. S. Packer et al., A lineage-resolved molecular atlas of C. elegans embryogenesis at single-cell resolution. Science 365 (2019).

33. T. J. Boyle, Z. Bao, J. I. Murray, C. L. Araya, R. H. Waterston, AceTree: a tool for visual analysis of Caenorhabditis elegans embryogenesis. BMC Bioinformatics 7, 275 (2006).

34. T. Walton et al., The Bicoid class homeodomain factors ceh-36/OTX and unc-30/PITX cooperate in C. elegans embryonic progenitor cells to regulate robust development. PLoS Genet 11, e1005003 (2015).

35. J. D. Rumley et al., pop-1/TCF, ref-2/ZIC and T-box factors regulate the development of anterior cells in the C. elegans embryo. Dev Biol 489, 34–46 (2022).

36. J. L. Richards, A. L. Zacharias, T. Walton, J. T. Burdick, J. I. Murray, A quantitative model of normal Caenorhabditis elegans embryogenesis and its disruption after stress. Dev Biol 374, 12–23 (2013).

37. E. Itskovits, R. Ruach, A. Kazakov, A. Zaslaver, Concerted pulsatile and graded neural dynamics enables efficient chemotaxis in C. elegans. Nat Commun 9, 2866 (2018).

38. T. Hino, S. Hirai, T. Ishihara, M. Fujiwara, EGL-4/PKG regulates the role of an interneuron in a chemotaxis circuit of C. elegans through mediating integration of sensory signals. Genes Cells 26, 411–425 (2021).

39. J. M. Gray, J. J. Hill, C. I. Bargmann, A circuit for navigation in Caenorhabditis elegans. Proc Natl Acad Sci U S A 102, 3184–3191 (2005).

40. A. Gordus, N. Pokala, S. Levy, S. W. Flavell, C. I. Bargmann, Feedback from network states generates variability in a probabilistic olfactory circuit. Cell 161, 215–227 (2015).

41. S. H. Chalasani et al., Neuropeptide feedback modifies odor-evoked dynamics in Caenorhabditis elegans olfactory neurons. Nat Neurosci 13, 615–621 (2010).

42. S. H. Chalasani et al., Dissecting a circuit for olfactory behaviour in Caenorhabditis elegans. Nature 450, 63–70 (2007).

43. C. I. Bargmann, E. Hartwieg, H. R. Horvitz, Odorant-selective genes and neurons mediate olfaction in C. elegans. Cell 74, 515–527 (1993).

44. Y. Sztainberg et al., Reversal of phenotypes in MECP2 duplication mice using genetic rescue or antisense oligonucleotides. Nature 528, 123–126 (2015).

45. E. Giacometti, S. Luikenhuis, C. Beard, R. Jaenisch, Partial rescue of MeCP2 deficiency by postnatal activation of MeCP2. Proc Natl Acad Sci U S A 104, 1931–1936 (2007).

46. S. K. Garg et al., Systemic delivery of MeCP2 rescues behavioral and cellular deficits in female mouse models of Rett syndrome. J Neurosci 33, 13612–13620 (2013).

47. H. T. Bjornsson et al., Histone deacetylase inhibition rescues structural and functional brain deficits in a mouse model of Kabuki syndrome. Sci Transl Med 6, 256ra135 (2014).

48. J. D. Rodriguez, D. J. Katz, Lineage Tracing and Single-Cell RNA-seq in C. elegans to Analyze Transgenerational Epigenetic Phenotypes Inherited from Germ Cells. Methods Mol Biol 2677, 61–79 (2023).

49. A. Santella, Z. Du, S. Nowotschin, A. K. Hadjantonakis, Z. Bao, A hybrid blob-slice model for accurate and efficient detection of fluorescence labeled nuclei in 3D. BMC Bioinformatics 11, 580 (2010).

50. J. I. Murray et al., The anterior Hox gene ceh-13 and elt-1/GATA activate the posterior Hox genes nob-1 and php-3 to specify posterior lineages in the C. elegans embryo. PLoS Genet 18, e1010187 (2022).

51. B. Katzman, D. Tang, A. Santella, Z. Bao, AceTree: a major update and case study in the long term maintenance of open-source scientific software. BMC Bioinformatics 19, 121 (2018).

52. J. Schindelin et al., Fiji: an open-source platform for biological-image analysis. Nat Methods 9, 676–682 (2012).

53. E. R. Santiago, A. Shelar, N. T. M. Christie, M. R. Lewis-Hayre, M. R. Koelle, Using NeuroPAL Multicolor Fluorescence Labeling to Identify Neurons in C. elegans. Curr Protoc 2, e610 (2022).

