## Supplemental Figures for "Abnormal behavior is reversible in a chromatin mutant"

**Supplementary figures**

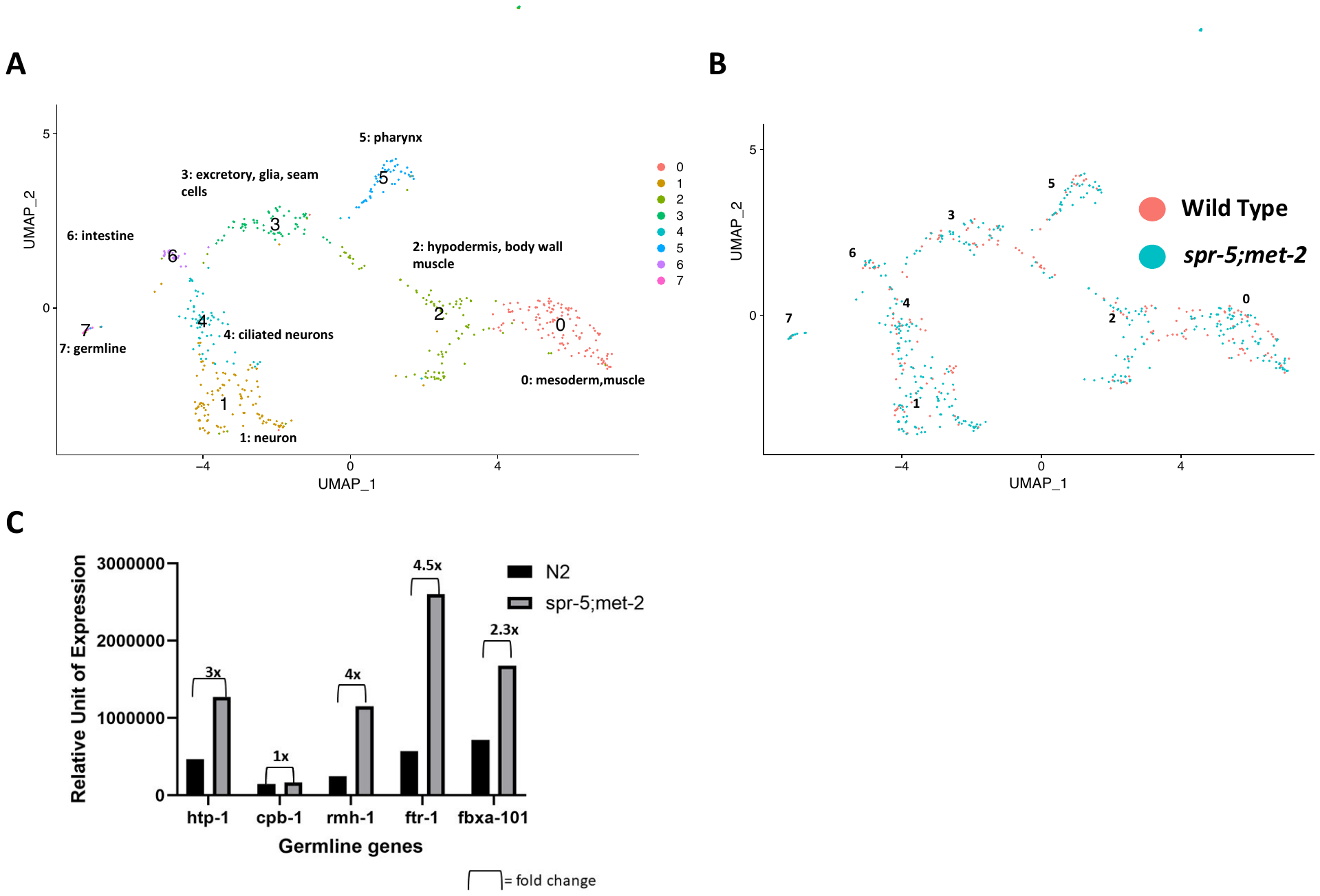

**Fig. S1. Germline genes are ectopically expressed, but overall gene expression is largely unchanged during embryogenesis in *spr-5; met-2* mutants.**

Unsupervised hierarchical clustering of single cell RNAseq data forms 8 clusters (**A**) and these clusters are the same between Wild Type (pink) and *spr-5; met-2* mutants (aqua) (**B**). (**C**) Quantitative RT-PCR data showing the relative gene expression of 5 MES-4 targeted germline genes not significantly misexpressed in our single-cell dataset but found to be significantly misexpressed at the L1 stage in *spr-5; met-2* mutants in our previous work (20). The data were normalized against the expression of *ama-1* from embryos at the 100-200 cell stage.

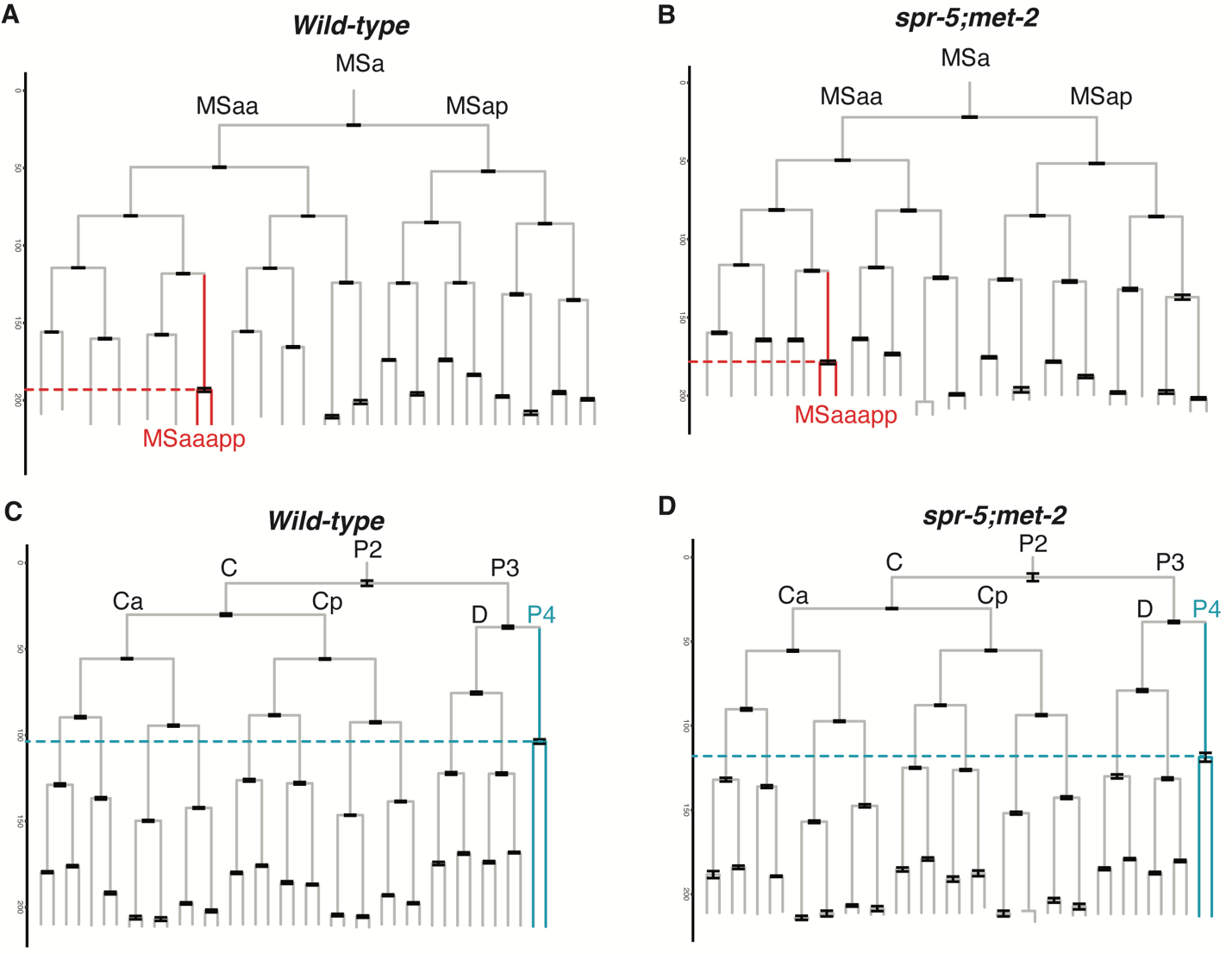

**Fig. S2**. **Comparison of the embryonic cell lineage in *spr-5; met-2* versus Wild Type.**

Wild-type (N2) (**A,C**) and *spr-5; met-2* (**B,D**) embryonic lineages highlighting the earlier MSaaapp cell division (red dashed boxes in Fig. 1A,B) and the later P4 cell division (blue dashed boxes in Fig. 1A,B) in *spr-5; met-2* compared to Wild Type. The error bars indicate the SEM from 22 wild-type and 8 *spr-5; met-2* lineages.

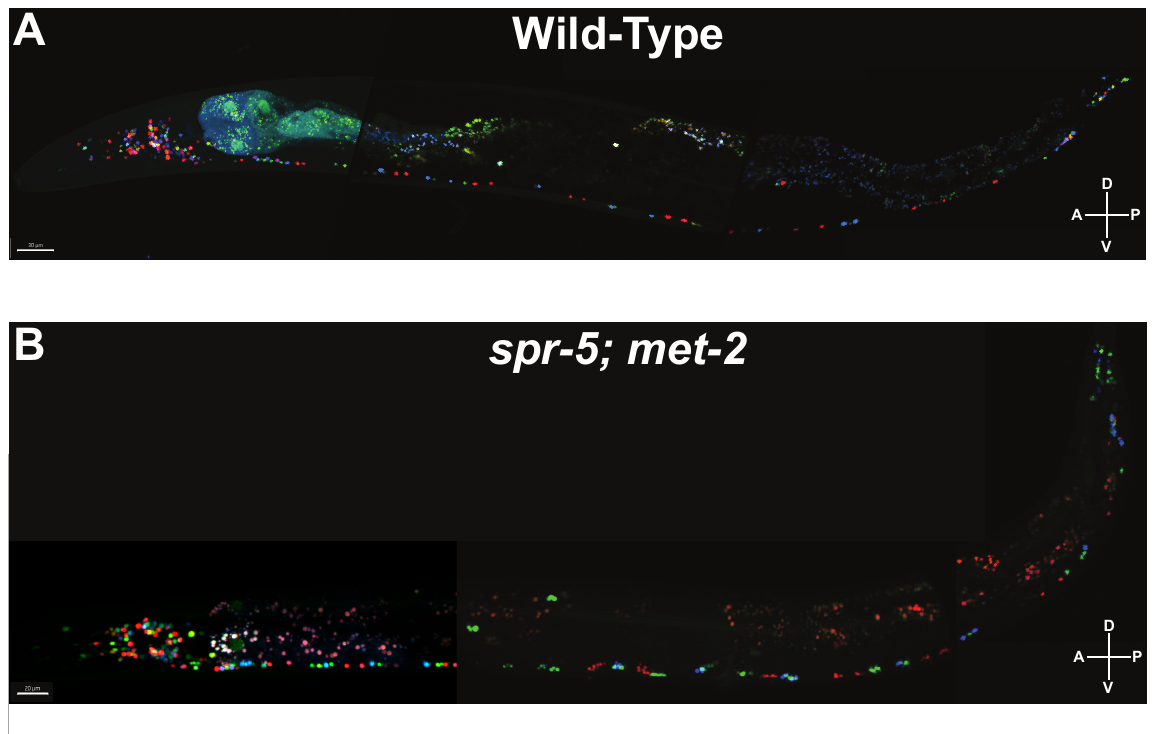

**Fig. S3. The nervous system is completely intact in *spr-5; met-2* mutants with impaired chemotaxis.**

Example of wild-type (**A**) and *spr-5; met-2* (**B**) adult worms indicating that all 302 uniquely identified neurons are present in the correct position. The anterior(A)/posterior(P) and dorsal(D)/ventral (V) axes are indicated.

**Supplementary Information**

Dataset S1 lineage tracing rate- corrected rate cycles (related to Fig. 2)

Dataset S2 lineage tracing- specific rates (related to Fig. 2)

Video S1A WT average projection (related to Fig. 3)

Video S1B *spr-5; met-2* average projection (related to Fig. 3)

Video S2A WT chemotaxis on OP50 bacteria (related to Fig. 3)

Video S2B *spr-5; met-2* chemotaxis on HT115 bacteria (related to Fig. 3)

Video S2C *spr-5; met-2* on *mes-4* RNAi chemotaxis (related to Fig. 3)

Video S2D WT chemotaxis on HT115 bacteria (related to Fig. 3)

Video S2E *spr-5; met-2* on *lsl-1* RNAi chemotaxis (related to Fig. 3)

Video S3A WT NeuroPAL rotation (related to Fig. 4)

Video S3B *spr-5; met-2* NeuroPAL rotation (related to Fig. 4)
